# Repeated footshock stress induces an escalation of cocaine self-administration in male and female rats: Role of the cannabinoid receptor 1

**DOI:** 10.1101/2023.02.23.529774

**Authors:** Andrew D. Gaulden, Erin A. Tepe, Eleni Sia, Sierra S. Rollins, Jayme R. McReynolds

**Affiliations:** Department of Pharmacology & Systems Physiology, University of Cincinnati College of Medicine, Cincinnati, OH; Center for Addiction Research, University of Cincinnati College of Medicine, Cincinnati, OH

**Author notes:** Corresponding Author: Jayme McReynolds, PhD, Address: 2120 E. Galbraith Rd #A-141, Cincinnati, OH 45237.

## Abstract

Stress is a significant contributor to the development and progression of substance use disorders (SUDs) and is problematic as it is unavoidable in daily life. Therefore, it is important to understand the neurobiological mechanisms that underlie the influence of stress on drug use. We have previously developed a model to examine the contribution of stress to drug-related behavior by administering a stressor, electric footshock stress, daily at the time of cocaine self-administration in rats resulting in an escalation of cocaine intake. This stress-induced escalation of cocaine intake involves neurobiological mediators of stress and reward such as cannabinoid signaling. However, all of this work has been conducted in male rats. Here we test the hypothesis that repeated daily stress can produce an escalation of cocaine in both male and female rats. We further hypothesize that cannabinoid receptor 1 (CB1R) signaling is recruited by repeated stress to influence cocaine intake in both male and female rats. Male and female Sprague-Dawley rats self-administered cocaine (0.5 mg/kg/inf, i.v.) during a modified short-access paradigm wherein the 2-hr access was separated into 4-30 min self-administration blocks separated by 4-5 min drug free period. Footshock stress produced a significant escalation of cocaine intake similarly in both male and female rats. Female stress-escalated rats did display greater time-out non-reinforced responding and greater “front-loading” behavior. In males, systemic administration of a CB1R inverse agonist/antagonist Rimonabant only attenuated cocaine intake in rats with a history of combined repeated stress and cocaine self-administration. However, in females, Rimonabant attenuated cocaine intake in the no stress control group but only at the highest dose of Rimonabant (3 mg/kg, i.p.) suggesting that females show a greater sensitivity to CB1R antagonism. However, female rats with a history of stress showed even greater sensitivity to CB1R antagonism as both doses of Rimonabant (1, 3 mg/kg) attenuated cocaine intake in stress-escalated rats similar to males. Altogether these data demonstrate that stress can produce significant changes in cocaine self-administration and suggests that repeated stress at the time of cocaine self-administration recruits CB1Rs to regulate cocaine-taking behavior across sexes.

## 1. Introduction

Cocaine use is a growing public health concern associated with serious harm to physical, interpersonal, and economic well-being. Currently, around 1.4 million individuals in the U.S. have cocaine use disorder (CUD) (SAMSHA 2022). To date, there is currently no FDA-approved treatment for CUD, and psychosocial interventions have limited efficacy and adherence (Kampman, 2019; Bentzley et al., 2021). CUD is a complex disorder with many factors that may influence continued cocaine use. Stress has been identified as a potential driver of cocaine use and craving for individuals with CUD. Indeed, experimentally delivered stressors increased cocaine craving for individuals who were cocaine-abstinent (Sinha and Li, 2007) and recent users (Kexel et al., 2022). Further, early life stress is a strong predictor for cocaine use disorder (Felitti, 2003; Elton et al., 2014; Spatz Widom et al., 1999; Rovaris et al., 2015), and CUD is strongly associated with stress-associated psychiatric illness (Chen et al., 2011). Individuals with substance use disorders have a high comorbidity with post-traumatic stress disorder (PTSD) (Back et al., 2006), and several studies indicate that individuals with PTSD use more cocaine in response to negative life events (Waldrop, Back, Brady, Upadhyaya, McRae and Saladin, 2007; Back et al., 2006; Waldrop, Back, Verduin et al., 2007). Negative life events can similarly drive cocaine use for the general population (Waldrop, Back, Brady, Upadhyaya, McRae and Saladin, 2007). Because stress is an unavoidable risk factor, and because individuals who use cocaine generally experience more daily stressors (Waldrop, Back, Brady, Upadhyaya, McRae and Saladin, 2007), it is important to understand the mechanisms by which stress contributes to cocaine use.

Studies using preclinical models of stress-cocaine interactions mostly examine the context of either acquisition of cocaine self-administration (SA) (Haney et al., 1995; Hofford et al., 2017; Tidey and Miczek, 1997) or reinstatement of cocaine-seeking behavior (Boutrel et al., 2005; Redila and Chavkin, 2008; Capriles et al., 2003; Mantsch and Goeders, 1999). However, less is known about how stressors can influence stable cocaine SA. We and others have previously shown that footshock stress, administered daily at the time of cocaine SA in the drug context, can escalate cocaine SA in otherwise stable responding male rats (Mantsch and Katz, 2006; McReynolds et al., 2022). This method for escalation is time- and context-dependent, and dependent on stress-evoked glucocorticoid release (Mantsch and Katz, 2006). Additionally, we have shown that this paradigm produces long-lasting changes in cocaine-motivated behavior. In rats with a history of repeated footshock stress, cocaine SA remains elevated following removal of the stressor, and cocaine-seeking behavior is enhanced long after the last stress exposure (McReynolds et al., 2022).

Preclinical literature investigating stress-cocaine interactions is also limited by male-only experiments. Despite this limitation, growing evidence suggests that females are more likely to escalate cocaine use (Roth and Carroll, 2004; Calipari et al., 2017; Becker and Hu, 2008). A larger body of preclinical data indicates that females show more autonomic activation to some stressful stimuli (Kudielka and Kirschbaum, 2005; Heck and Handa, 2018; Trainor, 2011) and have a higher prevalence of relevant stress-associated psychiatric illness (Bangasser and Valentino, 2014). Given the propensity of stress enhancement in cocaine SA and the reported sexual dimorphisms in cocaine- and stress-related behaviors, it is important to understand what differences, if any, exist between males and females for stress-induced escalation of cocaine SA.

Our current understanding of how repeated stress enhances cocaine SA involves neuroadaptations at the interface of reward and stress-related neurocircuitry (Koob, George and Kreek, 2007; Han et al., 2017). The endocannabinoid (eCB) system is a neurotransmitter system that regulates stress and reward in reward-related regions of the brain to promote homeostatic adaptations. Cocaine use regulates eCB signaling (Arnold, 2005; Parsons and Hurd, 2015), and polymorphisms in the cannabinoid type one receptor (CB1R) is associated with cocaine dependence (Clarke et al., 2013). In acute and chronic stress conditions, eCB levels increase (Morena et al., 2015), particularly in reward-related regions of the brain (Luján et al., 2021). Previous work has shown that under non-escalating, short-access cocaine SA conditions, systemic CB1R antagonism does not significantly attenuate cocaine SA (Orio et al., 2009; Bystrowska et al., 2018). However, under long-access cocaine SA conditions, which reliably produces escalation, systemic CB1R antagonism can reduce motivation for cocaine as measured by reduction in the breakpoint under a progressive ratio schedule of reinforcement. Additionally, we have demonstrated that systemic CB1R antagonism attenuates cocaine SA but only in rats with a history of combined stress and cocaine SA (McReynolds et al., 2022).

However, all of these data were collected in male rats only. Therefore, this work is designed to compare stress-induced escalation of cocaine SA in male and female rats, and the extent to which CB1R function regulates stress-escalated cocaine SA. We hypothesize that females will show more stress-induced escalation of cocaine SA, and that this effect is mediated by dimorphic differences in CB1R function.

## 2. Materials and Methods

### 2.1. Subjects

65 male and freely-cycling female rats (approx. 70 days old; Males: 250-275 g at arrival; Females: 225-250g at arrival; Envigo, Indianapolis, IN) were housed individually in a temperature- and humidity-controlled AAALAC-accredited animal facility on a reverse 12:12 hr light cycle (9 am lights off). Rats were given *ad libitum* access to food and water. Procedures were approved by the Institutional Animal Care and Use Committee at the University of Cincinnati and conducted in compliance with NIH guidelines. All experiments were conducted during the dark, active phase of the rat’s light cycle.

### 2.2. Surgery

All rats received implantation of intravenous catheters. Rats were anesthetized with isoflurane (2-2.5% in O_2_; Covetrus North America, Portland, ME) and the catheter was implanted into the superior vena cava with the backmount situated approx. 2-2.5 cm behind the rat’s scapula. The catheter consisted of a back-mounted cannula (Plastics One, Roanoke, VA) connected on the underside of the backmount to polypropylene mesh (500 microns; Small Parts, Logansport, IN). The backmount cannula was attached to a polyurethane catheter (0.6 mm i.d. x 1.1 mm o.d.; Access Technologies, Skokie, Il). Rats were given meloxicam (1mg/kg, sc) at the cessation of the surgery and for 2 additional days. The catheters were flushed with Cefazolin antiobiotic treatment (100 mg/kg, i.p.) in heparinized bacteriostatic 0.9% saline for 6 days following surgery. Following surgery recovery, the Cefazolin treatment ceased but the catheters continued to be flushed daily with heparinized bacteriostatic 0.9% saline throughout the experiment. Rats recovered for a minimum of one week before the initiation of self-administration.

### 2.3. Stress-induced escalation of cocaine

#### 2.3.1. Cocaine self-administration (SA)

All experiments were completed using Med Associates (Fairfax, VT) operant conditioning chambers in sound-attenuating cubicles. All operant chambers were interfaced with computers and procedures were executed and responding recorded using Med-PC V software. Each operant chamber contained two levers on the same wall with stimulus lights situated above the lever, a house light, and equipped for delivery of footshock through stainless steel grid floors. Rats initially were food deprived to approximately 90% of their starting weight and trained to lever press to receive a 45 mg sucrose pellet (Bio-Serv, Flemington, NJ) on a fixed ratio (FR) 1 schedule of reinforcement. Once rats successfully acquired lever pressing behavior, they were transitioned to delivery of a cocaine infusion (0.5 mg/kg/0.2 mL infusion, i.v.) in response to lever pressing using a modified short-access paradigm (Fig. 1). In this modified short-access paradigm rats were allowed to self-administer cocaine in 4 x 30 min SA blocks interspersed by 4 x 5 min drug-free periods. During the drug-free period the levers were retracted, the stimulus light was turned off, and the houselight turned on. During the SA block, the houselight turned off, levers were extended, and the stimulus light above the active lever was turned on. Once the rat received an infusion of cocaine, the active lever stimulus light turned off for a 10-sec period during which lever presses were recorded but not reinforced. Following the time-out period, the active lever stimulus light was re-illuminated and responding was once again reinforced. During the SA blocks, responding on a second, inactive lever was also recorded but not reinforced. Once rats showed regular responding on an FR1 schedule, they were gradually moved to an FR4 schedule for the remainder of the experiment. Stable baseline responding was defined as 3 days on FR4 with <10% variation across days. Once stable baseline responding was established rats were divided into the control no shock and shock stress groups balanced by baseline responding. The control footshock group continued to self-administer under the same conditions for 14 additional days. The shock stress group received unpredictable, intermittent electric footshock stress during the drug-free periods for 14 days.

**Figure 1.**
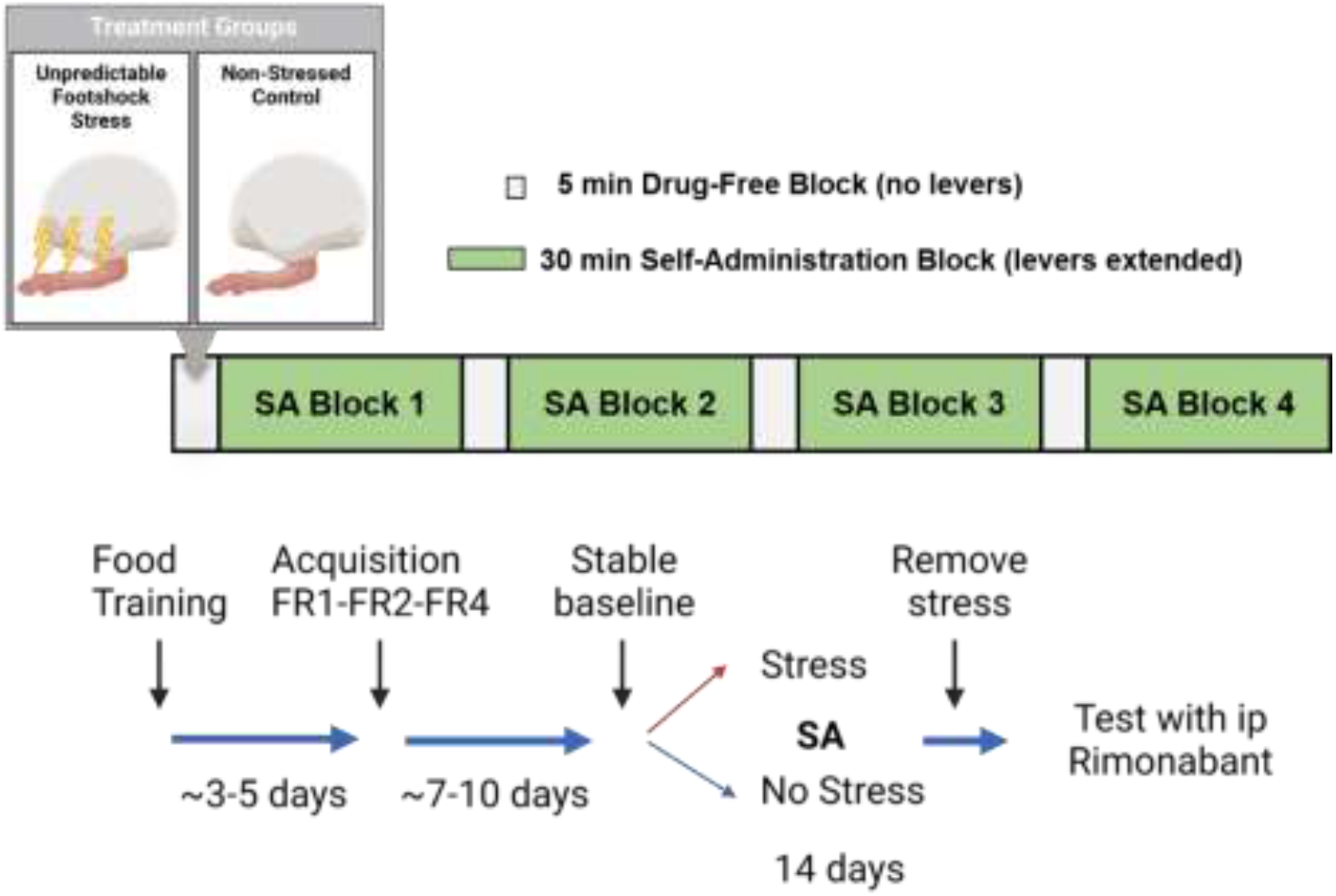
Experimental Design. A schematic depicting the modified short-access paradigm is on top. Briefly, the 2-hr short-access paradigm is divided into 4 x 30-min self-administration (SA) blocks. Separating these SA blocks are 4 x 5-min drug-free periods where the levers are retracted and different light cues are presented. After rats acquire stable lever pressing for cocaine on an FR4 schedule of reinforcement, half of the rats receive intermittent electric footshock stress during the drug-free period daily for 14 days. After 14 days of SA under stress or non-stress conditions, the footshock element is removed and rats can continue to SA cocaine. It is during this period that rats are tested for the ability of a CB1R inverse agonist/antagonist, Rimonabant, to influence cocaine SA.

#### 2.3.2. Footshock

Footshock administration consisted of triplets of shock (3 x 0.4 mA; 500 msec duration;

1 sec intershock interval) given at random intervals (30 sec average inter-triplet shock interval; range 15-60 sec). Rats received on average 7 triplet shock presentations per 5-min drug-free period for a total average of 28 triplet shocks for the whole self-administration session. The footshock was only administered during the drug-free period and was never administered when cocaine was freely available.

### 2.4. Effects of cannabinoid receptor 1 (CB1R) antagonism on cocaine-taking behavior

To test the role of CB1Rs in cocaine-taking behavior, some rats were pre-treated with a CB1R inverse agonist/antagonist prior to SA. Following 14 days of SA, footshock administration ceased and rats were allowed to continue cocaine SA and were given at least one day following cessation of the footshock before testing. The effects of the CB1R antagonist was tested under “no shock” conditions to isolate the contribution of CB1Rs to cocaine self-administration and eliminate any potential effects that would be related to the acute response to shock. On the testing day, rats received a systemic injection of Rimonabant (0, 1, 3 mg/kg, i.p.) 30-min prior to the beginning of the self-administration session. All rats received at least the vehicle and one dose of rimonabant in a counter-balanced order using a within-subject’s design. Most rats received all doses of Rimonabant. Any rat that did not receive a vehicle injection was excluded from study. There was at least one day of SA between tests to ensure the washout of the drug. Rats received a maximum of three tests total.

### 2.5. Drugs

Cocaine HCl and Rimonabant was obtained from the National Institute of Drug Abuse (NIDA) Drug Supply Program. Cocaine was dissolved in bacteriostatic 0.9% saline. Rimonabant was first dissolved in ethanol, followed by Cremaphor and finally by saline (0.9%) in a 1:1:18 ratio. The vehicle consisted of the 1:1:18 ratio of ethanol: Cremaphor: saline.

### 2.6. Data Analysis

Statistical analyses were conducted using GraphPad Prism (San Diego, CA). For comparison on sex differences in SA behavior, cocaine infusions and lever presses were analyzed in each sex separately using two-way repeated measures ANOVA or mixed-model repeated measures test (for any instances in which we were unable to collect all 14 days of SA data). To compare data across sexes, the infusion data was transformed to express each SA day as a percentage of baseline responding to account for differential baseline responding across sexes. These data were then analyzed using a three-way mixed model repeated measures ANOVA. For infusion data across SA 1-14, net area under the curve (AUC) was calculated for each animal with their average baseline responding set as zero and then AUCs across stress groups and sexes were analyzed using a two-way ANOVA. For Rimonabant testing, infusions and lever presses were analyzed using a two-way mixed-model repeated measures ANOVA. To assess the relationship between rimonabant-induced attenuation of cocaine SA and measures of SA, the rimonabant data was transformed to be represented as change from vehicle for each dose of rimonabant tested. A simple linear regression was used to test if measures of cocaine SA (AUC, cumulative infusions) could predicted rimonabant effects on cocaine SA. All ANOVAs were followed by Sidak or Dunnett’s post hoc testing where appropriate.

## 3. Results

### 3.1. Repeated footshock stress induces a significant escalation of cocaine self-administration in both male and female rats

Both male and female rats who received footshock stress at the time of cocaine self-administration show a significant escalation of cocaine SA (Fig. 2A-B). In males, a two-way mixed-model repeated measures test revealed a significant overall effect of SA day (Baseline (BL)-SA14; F(14,447)=2.07, p<.05) and a significant interaction between Stress condition and SA Day (F(14,447)=1.99, p<.05), though no overall effect of stress (No Shock vs Shock; F(1,32)=3.73, p=.06), on cocaine infusions across SA days (Fig. 2A). A Dunnett’s post hoc revealed significant differences across several days of SA compared to baseline but only in the Shock group (SA 2, 4-9, 11-12, p<.05), indicating that stress significantly escalated cocaine SA. In females, a two-way mixed-model repeated measures test revealed a significant overall effect of stress (No Shock vs Shock; F(1,29)=14.33, p<.001) and a significant interaction between Stress condition and SA Day (F(14,400)=1.92, p<.05), though no overall effect of SA Day (Baseline (BL)-SA14; F(14,400)=1.22, p=0.26), on cocaine infusions across SA days (Fig. 2B). A Dunnett’s post hoc revealed significant differences across several days of SA compared to baseline but only in the Shock group (SA 2-8, 11-14, p<.05), indicating that stress significantly escalated cocaine SA.

**Figure 2.**
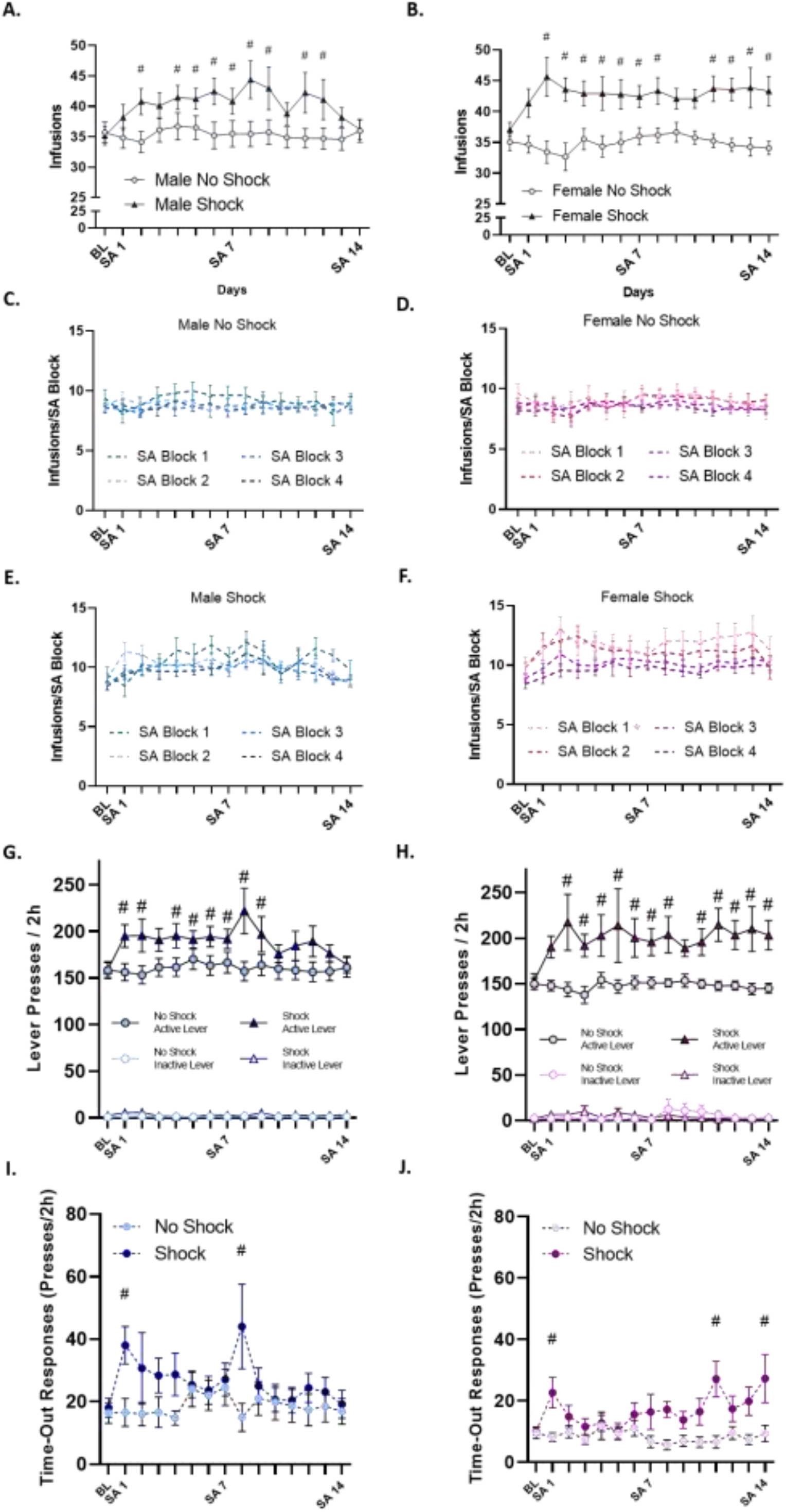
Repeated footshock stress induces a significant escalation of cocaine self-administration in both male (A) and female rats (B) as assessed by an increase in cocaine infusions across days. Male (C) and female (D) control non-stress rats show stable cocaine SA across the 4 SA blocks both within each day and across days. Male stress rats (E) show a significant increase in intake across SA days but cocaine SA is similar across SA blocks within each day. However, female stress rats (F) show a significant increase in cocaine intake across SA days and also show higher levels of cocaine SA in the first 2 SA blocks suggestive of increased frontloading behavior. Cocaine infusions in SA block 1 are significantly greater than SA Block 4 (*p<.05). Both male (G) and female (H) stress rats specifically show an increase in lever presses on the active reinforced lever and do not show differences in inactive lever pressing demonstrating that the increase in lever pressing is specific to cocaine. Time-out responding is defined as responding on the active lever during the time-out period immediately following a cocaine infusion that is recorded but not reinforced. While the male stress rats (I) show some increase in time-out responding, female stress rats (J) show significantly increased time-out responding across days that appears to be an emergent phenotype. (#p<.05, compared to baseline)

**Figure 3.**
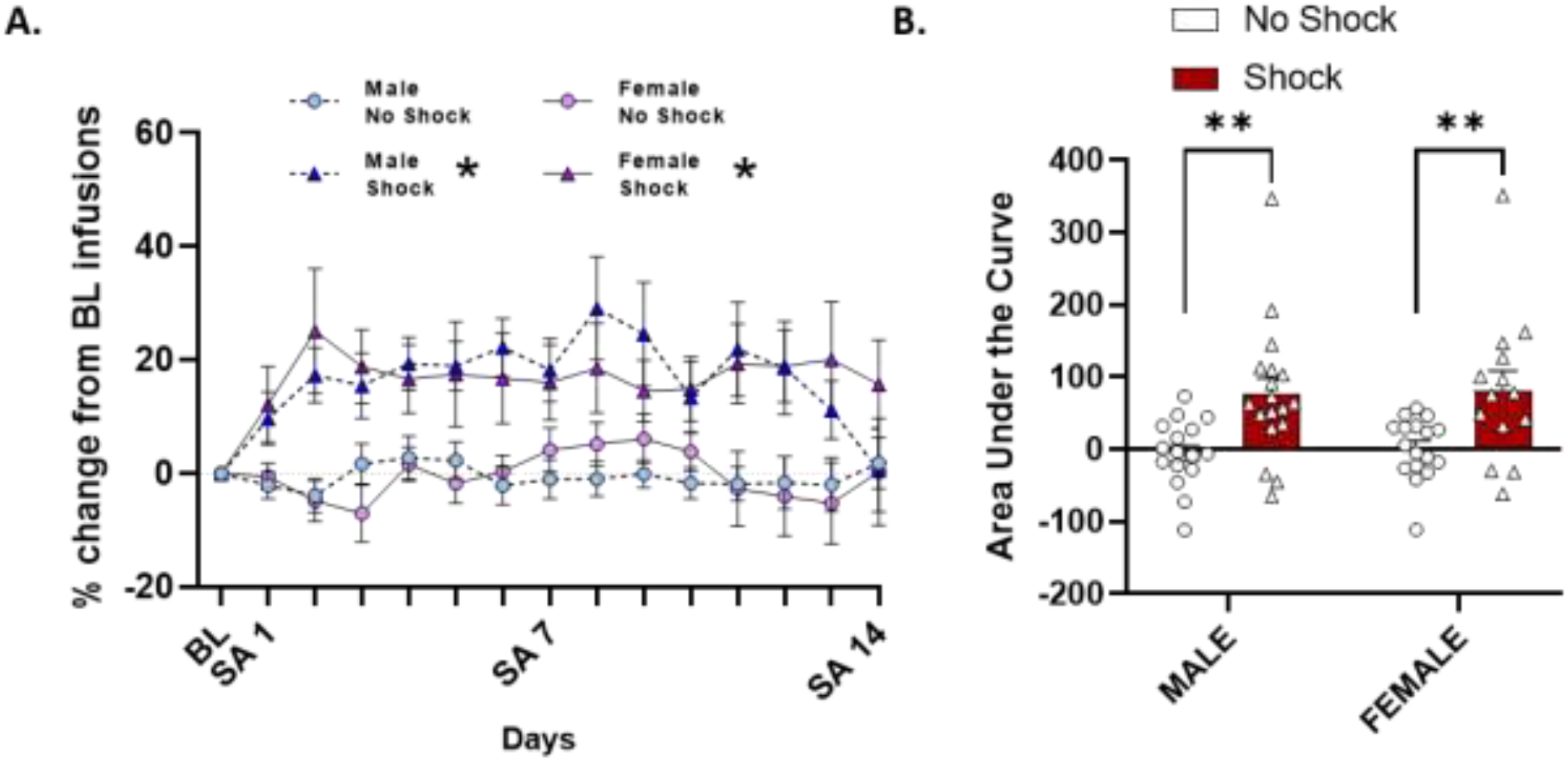
Male and female rats show similar levels of stress-induced escalation of cocaine SA. A) The infusion data was transformed to be represented each day as a percentage of baseline to account for difference in baseline infusions. While there is an overall stress and stress x SA day interaction (*p<.05) in both male and female rats, indicating that both sexes show a stress escalation phenotype, there is no effect of sex. Rates and levels of escalation are similar between male and female rats. B) Area under the curve was calculated for each rat with their baseline infusions set as zero. Rats in the stress group show significantly higher area under the curve measurements than the non-stress group though there is no significant difference between male and female rats.

**Figure 4.**
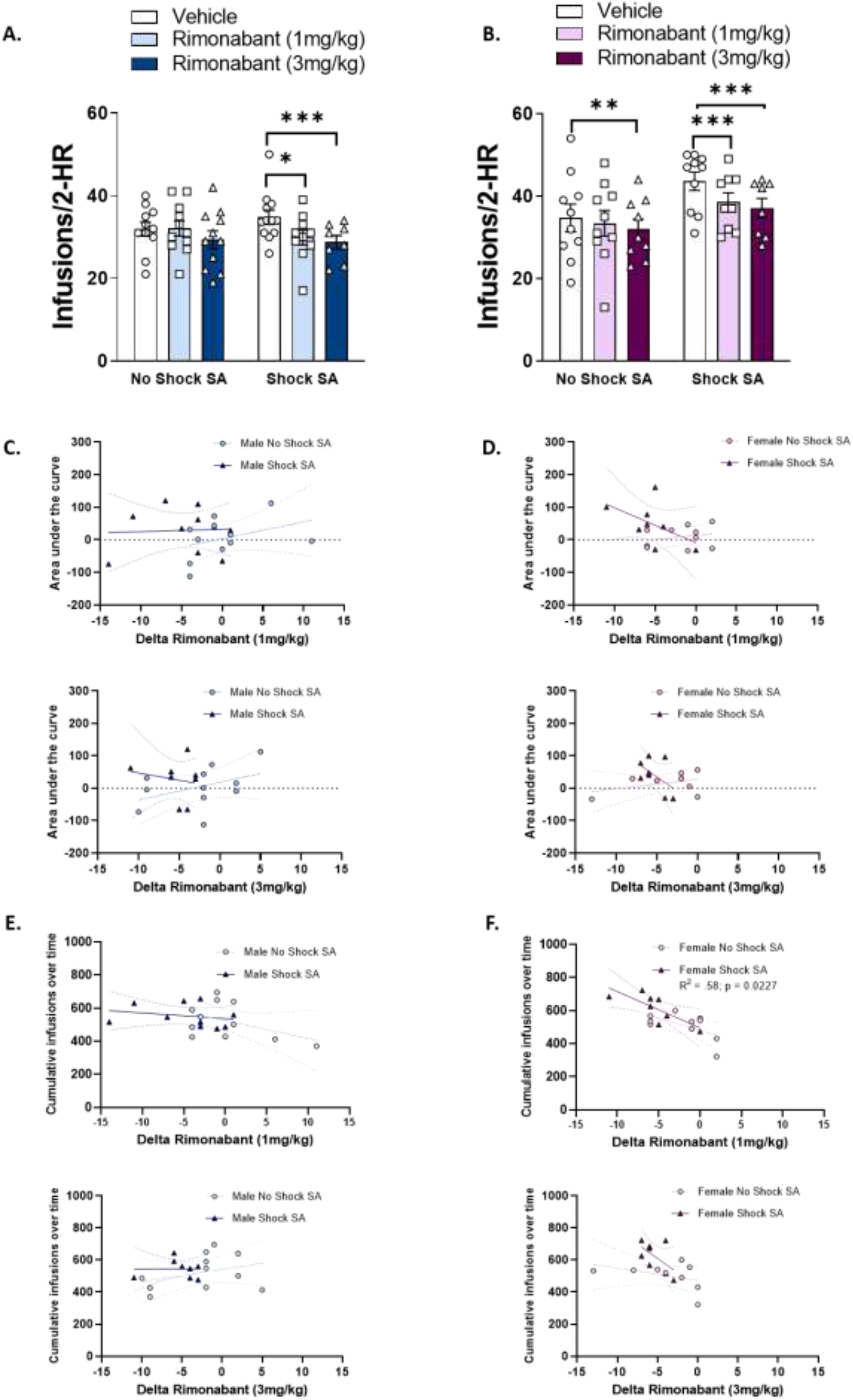
Systemic pre-treatment with CB1R antagonist Rimonabant 30 min prior to cocaine SA significantly attenuates cocaine intake in male (A) and female (B) rats with a prior history of stress at both doses tested (1, 3 mg/kg, i.p.). In the males, Rimonabant has no effect on cocaine intake in the non-stress rats but does significantly attenuate cocaine intake in the non-stress female rats at the highest dose (3 mg/kg, i.p.). The degree to which Rimonabant changed cocaine intake compared to vehicle (Delta Rimonabant) was assessed and a linear regression was used to determine whether measures of cocaine SA could predict changes in Delta Rimonabant. In both males (C) and female (D) rats, Delta Rimonabant was not associated with area under the curve in either stress or no stress rats. In male rats (E), Delta Rimonabant was not associated with cumulative infusions in either the stress or no stress group. However, in females, Delta Rimonabant at the 1 mg/kg dose was significantly correlated with cumulative infusions but only in the stress, and not non-stress, group.

We then examined whether cocaine SA differed across the 4 SA blocks within a day in the shock and no shock groups (Fig. 2C-F). In males, there was no significant difference in cocaine SA across blocks in either the no shock or shock groups. In the male no shock group (Fig. 2C), a twoway repeated measures ANOVA revealed no significant overall effect of SA Day (F (14, 840) = 1.162, p=0.30), SA Block (F (3, 60) = 0.31, p=0.82), or a significant interaction between SA Day and SA Block (F (42, 840) = 1.17, p=0.22). In the male shock group (Fig. 2E), a two-way mixed-model repeated measures test revealed no significant overall effect of SA Block (F (3, 68) = 0.72, p=0.54), and no significant interactions between SA Block and SA Day (F (42, 948) = 1.26, p=0.12), but an overall effect of SA Day (F (14, 948) = 5.72, p<.0001), indicating that while rats in the shock group showed significant changes in SA across days there were no significant changes in responding across SA blocks within each day demonstrating that the rats respond similarly within each block and don’t necessarily display “loading” behavior as it pertains to the 30-min SA blocks. In the females, the no shock group (Fig. 2D) is similar to the males, in that SA is similar across SA Blocks, as a two-way mixed model repeated measures test revealed no significant overall effect of SA Day (F (14, 876) = 1.61, p=0.07), no significant overall effect of SA Block (F (3, 64) = 0.45, p=0.72) and no significant interaction between SA Day and SA Block (F (42, 876) = 0.61, p=0.98). However, in the female shock group (Fig. 2F), there are differences in SA across SA blocks. A two-way repeated measures ANOVA revealed a significant overall effect of SA Block (F (3, 52) = 3.52, p<.05) and a significant overall effect of SA Day (F (14, 728) = 2.92, p<.001) but no significant interaction between SA Block and SA Day (F (42, 728) = 0.59, p=0.98). A Sidak’s post hoc test reveals that SA in SA Block 1 is significantly different from SA Block 4 (p<.05). This indicates that cocaine SA is greater in the first 30-min of SA than the last 30-min of SA and suggests that female rats exposed to stress engage in “front-loading” behavior that is not observed in either the female control no stress group or the male stress group.

Over the course of stress-induced escalation, both male and female rats show greater total responding on the active lever in the shock group as compared to the no shock group while not demonstrating increases in responding on the unreinforced inactive lever (Fig. 2G-H). In males (Fig. 2G), a two-way mixed-model repeated measures test revealed an overall effect of stress (F (1, 32) = 4.28, p<.05), and an overall effect of SA Day (F (14, 447) = 1.70, p=0.05), but no significant interaction between stress and SA Day (F (14, 447) = 1.68, p=.06) on total active lever presses across SA days. However, a two-way mixed model repeated measures test revealed no overall effect of stress (F (1, 32) = 3.10, p=0.09), overall effect of SA Day (F (14, 447) = 0.72, p=0.76), and no significant interaction between stress and SA Day (F (14, 447) = 0.50), on inactive lever presses across SA days. In females (Fig. 2H), a two-way mixed-model repeated measures test revealed a significant overall effect of stress (F (1, 29) = 10.01, p<.01), and a significant interaction between stress and SA Day (F (14, 401) = 1.71, p=0.05) but no overall effect of SA Day (F (14, 401) = 1.50, p=0.11) on total active lever presses across SA days. However, a two-way mixed model repeated measures test revealed no overall effect of stress (F (1, 29) = 0.01, p=0.93), no overall effect of SA Day (F (14, 401) = 0.72, p=0.76), and no interaction between stress and SA Day (F (14, 401) = 1.05, p=0.40) on inactive lever presses across SA days. Taken together, these data indicate that the increase in lever pressing in the Shock groups across sexes is specific to the active lever and not an overall increase in behavioral activation.

When examining total active lever presses, we noticed that lever presses in the female shock group continued to increase especially in the last several days of SA even though the total number of infusions did not continue to increase. Therefore, we isolated the time-out responding, responding on the active lever during the time-out period immediately following an infusion that is recorded but not reinforced, to assess whether there were any changes in time-out responding across SA days or as a result of the footshock stress (Fig. 2I-J). In the males (Fig. 2I), a two-way mixed model repeated measures test revealed no overall effect of SA Day (F (14, 447) = 0.99, p=0.46) or stress (F (1, 32) = 2.37, p=0.13) on time-out responding. While there was no significant interaction between SA Day and stress (F (14, 447) = 1.65, p=0.06) there was a trend. Planned comparisons with Dunnett’s post hoc testing revealed that males in the shock group had significantly greater time-out responses on SA1 and SA8 when compared to baseline (p<.05). This suggests that there were some acute stress-induced changes in time-out responding in males but that pattern of responding was not observed consistently across SA days. However, in the females (Fig. 2J), a two-way mixed model repeated measures test revealed a significant overall effect of SA Day (F (14, 387) = 1.87, p<.05), Stress (F (1, 28) = 10.13, p<.01), and an interaction between SA Day and Stress (F (14, 387) = 2.55, p<.01) on time-out responding. A Dunnett’s post hoc test revealed significant increases in time-out responding in the shock group on days SA1, SA11, and SA14 compared to baseline. In the female shock group there is an emergent increase in time-out responding across SA days suggesting potential increases in preservative or compulsive-like behavior.

### 3.2. Stress-induced escalation of cocaine self-administration is similar across sexes

Male and female rats had small differences in baseline cocaine SA, so to directly compare the effect of repeated footshock stress on cocaine SA across sexes, we transformed the infusion data for each SA day to be represented as a percentage change from baseline infusions. Then we analyzed both sexes together to examine any potential sex-specific difference in stress-induced escalation of cocaine intake across sexes (Fig. 2A). A three-way mixed model repeated measures test revealed while there is an overall effect of SA Day (F (14, 854) = 2.35, p<0.01), stress (F (1, 61) = 16.66, p<0.001), and an interaction between SA Day x stress (F (14, 854) = 2.60, p<0.01), there is no overall effect of Sex (F (1, 61) = 0.01, p=0.95), or an interaction between Stress x Sex (F (1, 61) = 0.001, p=0.97), SA Day x Sex (F (14, 854) = 0.50, p=0.93), or SA Day x Sex x Stress (F (14, 854) = 1.44, p=0.13). This indicates that cocaine SA in the no shock group is similar across sexes and that repeated footshock stress induces an escalation of cocaine intake in a similar manner across sexes as related to infusions across SA days. We also measured area under the curve (AUC) across SA 1-14 with baseline infusions for each rat serving as zero for the AUC measurement (Fig. 2B). A two-way ANOVA revealed a significant overall effect of stress (F (1, 61) = 18.35, p<0.0001), but no effect of Sex (F (1, 61) = 0.09, p=0.76) or a significant Sex x Stress interaction (F (1, 61) = 0.002, p=0.96) for AUC indicating that footshock stress escalates cocaine SA similarly between male and female rats.

### 3.3. Systemic administration of the CB1R inverse agonist/antagonist attenuates cocaine intake in stress-escalated male and female rats

T o test for the role of the CB1R in cocaine SA in rats with or without a history of stress, male and female rats received a systemic injection of the CB1R inverse agonist/antagonist, Rimonabant (1, 3 mg/kg, i.p.) or Vehicle 30 min prior to SA. This was done in a within-subjects design where rats received at least the vehicle and one dose of rimonabant and most rats received vehicle and both doses of rimonabant. In males, a two-way mixed model repeated measures test revealed a significant overall effect of the drug (F (2, 36) = 7.76, p<.01), but no overall effect of stress (F (1, 21) = 0.004, p=0.95) and no significant stress x drug interaction (F (2, 36) = 2.82, p=0.07). A Dunnett’s post hoc test revealed that Rimonabant at both doses significantly attenuated cocaine intake compared to Vehicle only in the stress group (1 mg, p<.01; 3 mg, p<.01) and not in the no stress group (1 mg, p=0.99; 3 mg, p=0.16). In the females, a two-way mixed model repeated measures test revealed a significant overall effect of the drug (F (2, 31) = 15.23, p<.0001), but no overall effect of stress (F (1, 18) = 3.74, p=0.07) and no significant stress x drug interaction (F (2, 31) = 2.30, p=0.12). A Dunnett’s post hoc tested revealed that in the no stress group, the 3 mg/kg dose of rimonabant significantly attenuated cocaine intake (p<.01) compared to the vehicle treatment, but not the 1 mg/kg dose (p=0.39). However, in the stress group, both doses of Rimonabant significantly attenuated cocaine intake (p<.001 for both doses) compared to vehicle treatment. This suggests that in both male and female rats, repeated stress is likely inducing neuroadaptations in the CB1R to influence cocaine SA. Furthermore, female rats may show some increased sensitivity to CB1R antagonism compared to males as we observe a significant effect of rimonabant in the no stress group at the highest dose.

We also assessed whether there was a relationship between measures of cocaine SA and the ability of rimonabant to attenuate cocaine intake. We first transformed the rimonabant data to be expressed as delta change from vehicle and used a linear regression test of whether the measures of cocaine SA, cumulative infusions or area under the curve, in the shock and no shock groups predicted rimonabant-induced attenuation of cocaine intake. In the males, the overall regression was not statistically significant for either the no shock (1 mg: R^2^=0.14, F(1,9)=1.52, p=0.25; 3 mg: R^2^= 0.18, F(1,9)=1.94, p=0.20) or the shock group (1 mg: R^2^=0.002, F(1,8)=0.02, p=0.89; 3 mg: R^2^=0.03, F(1,6)=0.22, p=0.66) for AUC at either dose (Fig. 2C). In males, the overall regression was also not statistically significant for either the no shock (1 mg: R^2^=0.21, F(1,9)=2.41, p=0.16; 3 mg: R^2^= 0.11, F(1,9)=1.16, p=0.31) or the shock group (1 mg: R^2^=0.06, F(1,8)=0.50, p=0.50; 3 mg: R^2^=0.001, F(1,6)=0.002, p=0.97) for cumulative infusions at either dose (Fig. 2E). In the females, the overall regression was not statistically significant for either the no shock (1 mg: R^2^=0.05, F(1,8)=0.38, p=0.55; 3 mg: R^2^= 0.17, F(1,6)=1.23, p=0.31) or the shock group (1 mg: R^2^=0.26, F(1,6)=2.12, p=0.20; 3 mg: R^2^=0.27, F(1,6)=2.19, p=0.19) for AUC at either dose (Fig. 2D). However, in females, while the overall regression was not statistically significant for the no shock group at either dose (1 mg: R^2^=0.36, F(1,8)=4.43, p=0.07; 3 mg: R^2^= 0.16, F(1,7)=1.31, p=0.29) or shock groups at the 3 mg dose (R^2^=0.33, F(1,6)=2.88, p=0.14), it was significant for the shock group at the 1 mg dose (R^2^=0.58, F(1,6)=8.35, p<.05) for cumulative infusions (Fig. 2F). This suggests that at the lower doses of rimonabant there may be a relationship because the extent of their escalation and intake and the effect of CB1R antagonism but interestingly only in the shock group.

## 4. DISCUSSION

Our experimental approach was designed to understand how a repeated footshock stress impacts cocaine SA for male and female rats, and how CB1R function influences cocaine SA under these conditions. This work revealed two key findings which improve our understanding of stress and CB1R involvement in cocaine SA for males and females. First, we found that our behavioral paradigm of administering a repeated footshock stress during cocaine SA produced a robust stress-induced escalation phenotype for both males and females. Additional analysis also revealed some key sex differences in the behavioral pattern of SA for stress-escalated male and female rats. The observed stress escalation phenotype for cocaine SA replicates our prior work (McReynolds et al., 2022), and extends the utility of this paradigm to female rats, which show a similar stress-induced escalation of cocaine SA. Based on our understanding that females are more likely to escalate cocaine usage than males (Becker and Hu, 2008; Roth and Carroll, 2004), we hypothesized that females would show more rapid and robust escalation compared to males. However, we did not observe clear sex differences in the escalation of cocaine intake for our stress SA groups. This may be attributed to our use of a short access SA paradigm, which does not produce escalation for males or females (Roth and Carroll, 2004). We also selected a weight-matched dose of cocaine that has been shown to produce stable SA responding in male and female rats, whereas a lower dose has been shown to produce female-specific escalation (Westenbroek et al., 2013). This work is limited by only providing one dose of cocaine, and future experiments using additional doses could yield more robust female-specific effects of stress on cocaine SA. Additionally, our stress paradigm is relatively novel; parameters such as shock frequency and/or intensity could be modified to leverage the elevated autonomic response to stressors seen in females given more established stress assays (Kudielka and Kirschbaum, 2005; Heck and Handa, 2018; Trainor, 2011; Doncheck et al., 2020). Although our paradigm does not fully model the sex differences in escalation of cocaine use seen in clinical literature, we do show subtle sex differences in SA behavior.

We demonstrate that within stress SA conditions, females alone show a unique “front-loading” behavior. This behavior has been previously shown in alcohol self-administration literature (Flores-Bonilla et al., 2021; Bauer et al., 2021), where females show greater consumption than males immediately after alcohol availability. The neurological basis for this behavior, and its sexually dimorphic regulation, remains unclear. Potential drivers of front loading in alcohol research include enhanced incentive salience for the drug after dopaminergic upregulation of reward circuitry (Ardinger et al., 2022) and front-loading to relieve negative symptoms from drug dependence (Koob, George F. and Le Moal, 2008; Ardinger et al., 2022). To our knowledge, very little is understood about front loading of cocaine self-administration, and how this behavior differs by sex. Further, we only see front loading in females who receive our stress-cocaine SA treatment. Whether our stressor acts as negative reinforcement to be relieved, or upregulates incentive salience neurocircuitry, or both, requires future investigation.

Our behavioral analysis also reveals the emergence of non-reinforced, “time-out” responding only for females in the stress SA group. These responses are *perseverative*; defined by continuous responses for the drug-associated stimulus when responses are not reinforced (Ersche et al., 2011). Perseverative responses are a type of compulsive-like behavior, which simply describes repetitive responses to an inappropriate situation, a key component of addiction (Dalley et al., 2011; Perry et al., 2008). Others have shown perseverative responding to a delay discounting food task after early life stress, but only for female rats (Brydges et al., 2015). Therefore, our behavioral paradigm provides evidence that females may be more susceptible to developing compulsive-like cocaine-motivated behavior after repeated stress. Understanding the mechanisms which drive this emergent phenotype may also characterize how females are more susceptible to psychostimulant addiction.

Second, our findings indicate that repeated stress in combination with cocaine SA causes long-term upregulation of CB1R signaling, and that females may be more sensitive overall to CB1R inactivation. Both male and female rats with a history of stress during cocaine self-administration showed significant attenuation of cocaine SA after pretreatment with the CB1R inverse agonist, Rimonabant. Importantly, while both doses of Rimonabant decreased cocaine infusions in male and female rats with a history of stress SA, non-stressed females showed sensitivity to the high dose of Rimonabant. This suggests that repeated stress during cocaine SA causes long-term changes in CB1R function, and that females without a history of stress during cocaine SA have additional sensitivity to CB1R inactivation that is not observed in males. CB1R activity may contribute to cocaine SA behavior via eCB signaling in components of the mesolimbic dopamine system, such as the ventral tegmental area (VTA) (Wang et al., 2015). Our recent work suggests that the long-term escalation from footshock stress during cocaine SA can be localized in part to changes in CB1R function, but not expression, at the VTA and nucleus accumbens (McReynolds et al., 2022). The precise mechanism by which stress and cocaine SA change eCB system function in reward circuitry remains unknown and is an ongoing research project of this group.

Rimonabant treatment revealed an effect of CB1R inactivation on cocaine SA for non-stressed females, but no significant effect for non-stressed males. This finding supports prior data (Orio et al., 2009; Bystrowska et al., 2018) showing that males do not show CB1R-dependent changes in cocaine SA under short-access, non-escalating conditions. The observed increase in sensitivity to CB1R inactivation in female non-stressed controls provides valuable information about sex-specific differences in eCB system function. To date, the bulk of research investigating sex effects in the eCB system focuses on conditions with robust neuroplasticity, such as stress effects on hippocampal LTP (Zer-Aviv and Akirav, 2016; Ferraro et al., 2020), and amygdala and prefrontal cortex eCB content (Vecchiarelli et al., 2022). Less is known about how cocaine experience might change eCB function for males and females, but recent studies suggest that female gonadal hormones, such as estradiol, modifies the molecular components of CB1R signaling within the nucleus accumbens (Peterson et al., 2016), a key structure for drug-motivated behavior. Although we did not track estrous staging for this set of experiments, understanding how estrous cycle and cocaine experience impacts female CB1R function remains an area of interest. Additional analysis revealed that females with a history of shock during cocaine SA had a significant correlation between the degree of rimonabant-mediated attenuation of cocaine SA and total cumulative intake of cocaine across SA days. We interpret this finding as a representation of long-lasting changes in stress and cocaine-evoked neuroplasticity (Mantsch et al., 2014) that may be more robust from augmented female stress responsivity (Heck and Handa, 2018; Trainor, 2011; Kudielka and Kirschbaum, 2005).

Together, these data suggest that concurrent footshock stress during cocaine SA produces escalation of cocaine intake in males and females under otherwise stable SA conditions. This escalation likely causes long-term changes in CB1R function, such that CB1R inactivation has a greater effect on attenuating cocaine intake for rats with a history of stress during SA. We also provide behavioral data showing more perseverative behavior only in females given stress during SA, which provides additional framework for how understanding how stress may contribute to the development of compulsive-like behavior in females. Stress is a pervasive and unavoidable risk factor for individuals who use cocaine. Therefore, understanding how stressors may contribute to the development and expression of cocaine use is critical for a more complete understanding of CUD. Additionally, a detailed understanding of how biological mediators such as the eCB system and biological sex may regulate stress and cocaine use will provide new perspectives and treatment options for treating CUD.

## 5. Acknowledgements

This work was supported by a National Institute on Drug Abuse Grant K01-DA045295 to JRM.

## Notes

### Competing Interest Statement

The authors have declared no competing interest.

## References

Substance abuse and mental health services administration (SAMHSA) (2022). Key substance use and mental health indicators in the united states: Results from the 2021 national survey on drug use and health (HHS publication no. PEP22-07-01-005, NSDUH series H-57). Center for Behavioral Health Statistics and Quality, Substance Abuse and Mental Health Services Administration.

Ardinger CE, Lapish CC, Czachowski CL, Grahame NJ (2022) A critical review of front-loading: A maladaptive drinking pattern driven by alcohol’s rewarding effects. Alcohol Clin Exp Res (HOBOKEN) 46:1772–1782.

Arnold JC (2005) The role of endocannabinoid transmission in cocaine addiction. Pharmacology, Biochemistry and Behavior (OXFORD) 81:396–406.

Back SE, Brady KT, Jaanimägi U, Jackson JL (2006) Cocaine dependence and PTSD: A pilot study of symptom interplay and treatment preferences. Addict Behav (OXFORD) 31:351–354.

Bangasser DA, Valentino RJ (2014) Sex differences in stress-related psychiatric disorders: Neurobiological perspectives. Front Neuroendocrinol (SAN DIEGO) 35:303–319.

Bauer MR, McVey MM, Boehm SL (2021) Three weeks of binge alcohol drinking generates increased alcohol Front-Loading and robust Compulsive-Like alcohol drinking in male and female C57BL/6J mice. Alcohol Clin Exp Res (HOBOKEN) 45:650–660.

Becker JB, Hu M (2008) Sex differences in drug abuse. Front Neuroendocrinol (SAN DIEGO) 29:36–47.

Bentzley BS, Han SS, Neuner S, Humphreys K, Kampman KM, Halpern CH (2021) Comparison of treatments for cocaine use disorder among adults: A systematic review and meta-analysis. JAMA Network Open (CHICAGO) 4:e218049.

Boutrel B, Kenny PJ, Specio SE, Martin-Fardon R, Markou A, Koob GF, de Lecea L (2005) Role for hypocretin in mediating stress-induced reinstatement of cocaine-seeking behavior. Proceedings of the National Academy of Sciences - PNAS (WASHINGTON) 102:19168–19173.

Brydges NM, Holmes MC, Harris AP, Cardinal RN, Hall J (2015) Early life stress produces compulsive-like, but not impulsive, behavior in females. Behav Neurosci (WASHINGTON) 129:300–308.

Bystrowska B, Frankowska M, Smaga I, Pomierny-Chamioło L, Filip M (2018) Effects of cocaine self-administration and its extinction on the rat brain cannabinoid CB1 and CB2 receptors. Neurotoxicity Research (New York) 34:547–558.

Calipari E, Juarez B, Morel C, Walker D, Cahill M, Ramakrishnan C, Deisseroth K, Han M, Nestler E (2017) Estrous cycle-dependent alterations in cocaine affinity at the dopamine transporter underlie enhanced cocaine reward in females. Biological Psychiatry (1969) 81:S276.

Capriles N, Rodaros D, Sorge RE, Stewart J (2003) A role for the prefrontal cortex in stress- and cocaine-induced reinstatement of cocaine seeking in rats. Psychopharmacology (Berl) (NEW YORK) 168:66–74.

Chen KW, Banducci AN, Guller L, Macatee RJ, Lavelle A, Daughters SB, Lejuez CW (2011) An examination of psychiatric comorbidities as a function of gender and substance type within an inpatient substance use treatment program. Drug Alcohol Depend (CLARE) 118:92–99.

Clarke T, Bloch PJ, Ambrose-Lanci LM, Ferraro TN, Berrettini WH, Kampman KM, Dackis CA, Pettinati HM, O’Brien CP, Oslin DW, Lohoff FW (2013) Further evidence for association of polymorphisms in the CNR1 gene with cocaine addiction: Confirmation in an independent sample and meta-analysis. Addict Biol (Oxford, UK) 18:702–708.

Dalley JW, Everitt BJ, Robbins TW (2011) Impulsivity, compulsivity, and top-down cognitive control. Neuron (Cambridge, Mass.) (CAMBRIDGE) 69:680–694.

Doncheck EM, Liddiard GT, Konrath CD, Liu X, Yu L, Urbanik LA, Herbst MR, DeBaker MC, Raddatz N, Van Newenhizen EC, Mathy J, Gilmartin MR, Liu Q, Hillard CJ, Mantsch JR (2020) Sex, stress, and prefrontal cortex: Influence of biological sex on stress-promoted cocaine seeking. Neuropsychopharmacology (New York, N.Y.) (LONDON) 45:1974–1985.

Elton A, Smitherman S, Young J, Kilts CD (2014) Effects of childhood maltreatment on the neural correlates of stress- and drug cue-induced cocaine craving. Addict Biol (HOBOKEN) 20:820–831.

Ersche KD, Roiser JP, Abbott S, Craig KJ, Müller U, Suckling J, Ooi C, Shabbir SS, Clark L, Sahakian BJ, Fineberg NA, Merlo-Pich EV, Robbins TW, Bullmore ET (2011) Response perseveration in stimulant dependence is associated with striatal dysfunction and can be ameliorated by a D2/3 receptor agonist. Biological Psychiatry (1969) (New York, NY) 70:754–762.

Felitti VJ (2003) Origins of addictive behavior: Evidence from a study of stressful chilhood experiences. Prax Kinderpsychol Kinderpsychiatr (Germany) 52:547–559.

Flores-Bonilla A, De Oliveira B, Silva-Gotay A, Lucier KW, Richardson HN (2021) Shortening time for access to alcohol drives up front-loading behavior, bringing consumption in male rats to the level of females. Biology of Sex Differences (LONDON) 12:51.

Han X, DeBold JF, Miczek KA (2017) Prevention and reversal of social stress-escalated cocaine self-administration in mice by intra-VTA CRFR1 antagonism. Psychopharmacology (Berl) (Berlin/Heidelberg) 234:2813–2821.

Haney M, Maccari S, Le Moal M, Simon H, Vincenzo Piazza P (1995) Social stress increases the acquisition of cocaine self-administration in male and female rats. Brain Res (AMSTERDAM) 698:46–52.

Heck AL, Handa RJ (2018) Sex differences in the hypothalamic–pituitary–adrenal axis’ response to stress: An important role for gonadal hormones. Neuropsychopharmacology (New York, N.Y.) (LONDON) 44:45–58.

Hofford RS, Prendergast MA, Bardo MT (2017) A modified single prolonged stress episode delays acquisition of cocaine self-administration. Drug Alcohol Depend 171:e90.

Kampman KM (2019) The treatment of cocaine use disorder. Science Advances (WASHINGTON) 5:eaax1532.

Kexel A, Kluwe-Schiavon B, Baumgartner MR, Engeli EJE, Visentini M, Kirschbaum C, Seifritz E, Ditzen B, Soravia LM, Quednow BB (2022) Cue-induced cocaine craving enhances psychosocial stress and vice versa in chronic cocaine users. Translational Psychiatry (LONDON) 12:443.

Koob G, Kreek MJ (2007) Stress, dysregulation of drug reward pathways, and the transition to drug dependence. Am J Psychiatry (ARLINGTON) 164:1149–1159.

Koob GF, Le Moal M (2008) Addiction and the brain antireward system. Annual Review of Psychology (PALO ALTO) 59:29–53.

Kudielka BM, Kirschbaum C (2005) Sex differences in HPA axis responses to stress: A review. Biol Psychol (AMSTERDAM) 69:113–132.

Luján MÁ, Cheer JF, Melis M (2021) Choosing the right drug: Status and future of endocannabinoid research for the prevention of drug-seeking reinstatement. Current Opinion in Pharmacology (OXFORD) 56:29–38.

Mantsch JR, Katz ES (2006) Elevation of glucocorticoids is necessary but not sufficient for the escalation of cocaine self-administration by chronic electric footshock stress in rats. Neuropsychopharmacology (New York, N.Y.) (LONDON) 32:367–376.

Mantsch JR, Goeders NE (1999) Ketoconazole blocks the stress-induced reinstatement of cocaine-seeking behavior in rats: Relationship to the discriminative stimulus effects of cocaine. Psychopharmacology (Berl) (NEW YORK) 142:399–407.

Mantsch JR, Vranjkovic O, Twining RC, Gasser PJ, Mcreynolds JR, Blacktop JM (2014) Neurobiological mechanisms that contribute to stress-related cocaine use. Neuropharmacology 76:383–394.

McReynolds JR, Wolf CP, Starck DM, Mathy JC, Schaps R, Krause LA, Hillard CJ, Mantsch JR (2022) Role of mesolimbic endocannabinoid signaling in stress-driven cocaine use in rats. bioRxiv 2022.10.28.514315.

Morena M, Patel S, Bains JS, Hill MN (2015) Neurobiological interactions between stress and the endocannabinoid system. Neuropsychopharmacology (New York, N.Y.) (LONDON) 41:80–102.

Orio L, Edwards S, George O, Parsons LH, Koob GF (2009) A role for the endocannabinoid system in the increased motivation for cocaine in extended-access conditions. The Journal of Neuroscience (WASHINGTON) 29:4846–4857.

Parsons LH, Hurd YL (2015) Endocannabinoid signalling in reward and addiction. Nature Reviews.Neuroscience (BERLIN) 16:579–594.

Perry JL, Nelson SE, Carroll ME (2008) Impulsive choice as a predictor of acquisition of IV cocaine self-administration and reinstatement of cocaine-seeking behavior in male and female rats. Exp Clin Psychopharmacol (WASHINGTON) 16:165–177.

Redila VA, Chavkin C (2008) Stress-induced reinstatement of cocaine seeking is mediated by the kappa opioid system. Psychopharmacology (Berl) (Berlin/Heidelberg) 200:59–70.

Roth ME, Carroll ME (2004) Sex differences in the escalation of intravenous cocaine intake following long- or short-access to cocaine self-administration. Pharmacology, Biochemistry and Behavior (OXFORD) 78:199–207.

Rovaris DL, Mota NR, Bertuzzi GP, Aroche AP, Callegari-Jacques SM, Guimarães LSP, Pezzi JC, Viola TW, Bau CHD, Grassi-Oliveira R (2015) Corticosteroid receptor genes and childhood neglect influence susceptibility to crack/cocaine addiction and response to detoxification treatment. J Psychiatr Res (OXFORD) 68:83–90.

Sinha R, Li CSR (2007) Imaging stress- and cue-induced drug and alcohol craving: Association with relapse and clinical implications. Drug Alcohol Rev (Oxford, UK) 26:25–31.

Spatz Widom C, Weiler BL, Cottler LB (1999) Childhood victimization and drug abuse: A comparison of prospective and retrospective findings. J Consult Clin Psychol (WASHINGTON) 67:867–880.

Tidey JW, Miczek KA (1997) Acquisition of cocaine self-administration after social stress: Role of accumbens dopamine. Psychopharmacology (Berl) (NEW YORK) 130:203–212.

Trainor BC (2011) Stress responses and the mesolimbic dopamine system: Social contexts and sex differences. Horm Behav (SAN DIEGO) 60:457–469.

Waldrop AE, Back SE, Verduin ML, Brady KT (2007) Triggers for cocaine and alcohol use in the presence and absence of posttraumatic stress disorder. Addict Behav (OXFORD) 32:634–639.

Waldrop AE, Back SE, Brady KT, Upadhyaya HP, McRae AL, Saladin ME (2007) Daily stressor sensitivity, abuse effects, and cocaine use in cocaine dependence. Addict Behav (OXFORD) 32:3015–3025.

Wang H, Treadway T, Covey DP, Cheer JF, Lupica CR (2015) Cocaine-induced endocannabinoid mobilization in the ventral tegmental area. Cell Reports (Cambridge) (CAMBRIDGE) 12:1997–2008.

Westenbroek C, Perry AN, Becker JB (2013) Pair housing differentially affects motivation to self-administer cocaine in male and female rats. Behav Brain Res (AMSTERDAM) 252:68–71.

